# Unreliable homeostatic action potential broadening in cultured dissociated neurons

**DOI:** 10.1101/2025.05.09.653135

**Authors:** Andreas Ritzau-Jost, Salil Rajayer, Jana Nerlich, Filip Maciag, Alexandra John, Michael Russier, Victoria Gonzalez Sabater, Luke J. Steiger, Jacques-Olivier Coq, Jens Eilers, Maren Engelhardt, Juan Burrone, Dominique Debanne, Martin Heine, Stephen M Smith, Stefan Hallermann

## Abstract

Homeostatic plasticity preserves neuronal activity against perturbations. Recently, somatic action potential broadening was proposed as a key homeostatic adaptation to chronic inactivity in neocortical neurons. Since action potential shape critically controls calcium entry and neuronal function, broadening provides an attractive homeostatic feedback mechanism to regulate activity. Here, we report that chronic inactivity induced by sodium channel block does not broaden action potentials in neocortical neurons under a wide range of conditions. In contrast, action potentials were broadened in CA3 neurons of organotypic hippocampal cultures by chronic sodium channel block and in hippocampal dissociated cultures by chronic synaptic block. Mechanistically, BK-type potassium channels were proposed to underly inactivity-induced action potential broadening. However, BK channels did not affect action potential duration in our recordings. Our results indicate that action potential broadening can occur in specific neurons and conditions but is not a general mechanism of homeostatic plasticity in cultured neurons.

## Introduction

Action potentials (APs) control various functions including neurotransmitter release from axonal boutons^1^, calcium signals in the soma^2^, and synaptic plasticity in dendrites^3^. AP broadening during short-term plasticity on the time scale of milliseconds to seconds is well-established and occurs in many types of neuronal cell bodies^4,5^ and nerve terminals^6,7^. There are also indications that APs change during long-term plasticity on the time scale of minutes to hours. When examining Hebbian long-term plasticity (i.e., positive feedback-based and input-specific long-term plasticity^8^), some studies reported AP broadening. For example, presynaptic APs of hippocampal mossy fiber boutons broaden during depolarization-induced potentiation^9^ and somatic APs of neurons in the amygdala broaden during fear conditioning^10^. In contrast, the role of the AP duration is less clear for non-Hebbian homeostatic long-term plasticity (i.e., negative feedback-based and non-input specific long-term plasticity^8^). Specifically, homeostatic plasticity induced by neuronal inactivity in the presence of the sodium channel blocker tetrodotoxin (TTX) has been studied for decades, but AP broadening was initially not described^11,12^. A recent study however reports prominent AP broadening during TTX-induced homeostatic plasticity at the soma of cultured neocortical neurons^13^, which was not observed at presynaptic boutons of cultured neocortical neurons^14^. Compartmentalized AP broadening at the neuronal cell body would provide a strikingly simple and previously overlooked mechanism by which neuronal inactivity could increase subsequent excitability^13^. This mechanism involves decreased calcium entry during TTX application causing changes in gene expression that would result in broadened APs and more calcium entry after TTX removal. In this model, AP broadening is a key link in the homeostatic feedback loop underlying homeostatic plasticity. In a collaborative effort across several laboratories, we therefore systematically investigated AP broadening in cultured neurons during homeostatic plasticity under various experimental conditions.

## Results

### Cell type- and model-specific AP broadening in hippocampal neurons

APs in CA3 neurons of hippocampal organotypic slice cultures broaden homeostatically upon chronic inactivity using both, synaptic block by glutamate receptor antagonists^15^ or sodium channel block by TTX^16^ (see also ref. 17 for a similar trend). We first aimed to confirm homeostatic AP broadening in CA3 neurons in organotypic slice cultures. APs were elicited by depolarizing currents from the resting membrane potential during perfusion with TTX-free artificial cerebrospinal fluid. APs recorded in CA3 pyramidal neurons treated with TTX (48 hours) had longer durations than those in control cells (Fig. 1A, B; median [IQR] 1.75 [1.69–1.86] ms and 2.13 [2.07–2.22] ms for control and TTX-treatment, respectively, 14 cells each, P < 0.001), confirming earlier reports of homeostatic AP broadening in these neurons^15,16^.

**Figure 1.**
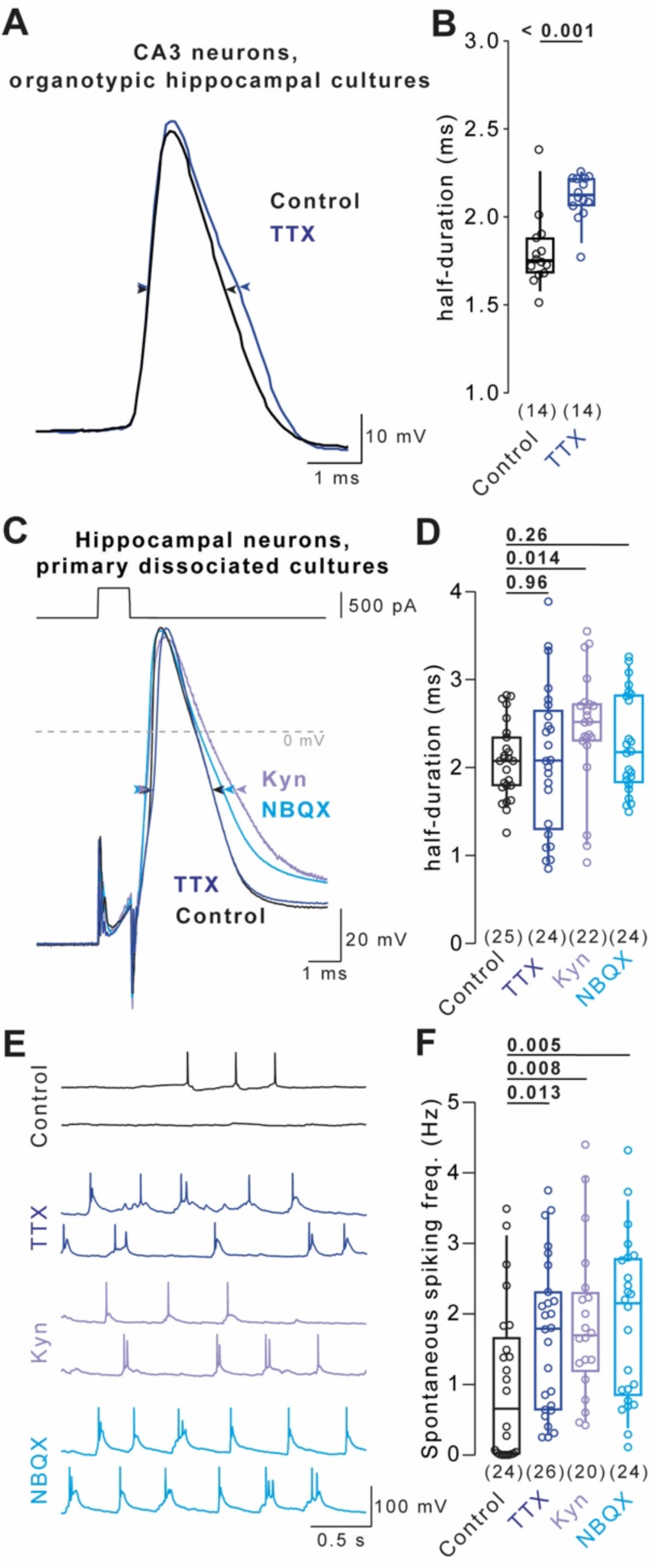
Cell type- and model-specific AP broadening in hippocampal neurons. (A) Example evoked APs recorded from CA3 pyramidal neurons in organotypic hippocampal cultures under control condition (black) and following 48 hours of TTX-treatment (blue). (B) AP durations recorded under conditions in A. (C) Example depolarization-evoked APs (500 pA for 1 ms) recorded from neurons in primary dissociated hippocampal cultures under control condition (black) and following 48 hours of TTX- (blue), Kyn- (purple), or NBQX-treatment (turquoise). (D) AP durations recorded under conditions in C. (E) Example spontaneous APs recorded under the conditions in C. (F) Spontaneous AP frequency under the conditions in C. Numbers in brackets reflect number of recorded neurons. Box plots as median ± interquartile range, whiskers indicate 10–90 percentile. P-values above graphs, significance tested by Mann-Whitney-*U* test.

Next, we tested whether homeostatic AP broadening occurred in dissociated hippocampal cultured neurons, a model widely used to study homeostatic plasticity. We inhibited neuronal activity for 48 hours to induce homeostatic plasticity by either blocking AP firing using TTX (Fig. 1C) or by blocking synaptic transmission using the low-affinity glutamate receptor blocker kynurenic acid (Kyn) or the high-affinity glutamate receptor blocker 2,3-Dioxo-6-nitro-1,2,3,4-tetrahydrobenzo [f]quinoxaline-7-sulfonamide (NBQX). Surprisingly, AP duration was unaffected by TTX-treatment in primary hippocampal cultured neurons (Fig. 1D; median [IQR] 2.07 [1.79–2.34] ms and 2.08 [1.32–2.61] ms for control and TTX, 25 and 24 cells, respectively, P = 0.96). In contrast, Kyn-treatment significantly increased AP duration in the same set of interleaved experiments (Fig. 1D; median [IQR] 2.52 [2.31–2.72] ms, 22 cells, P = 0.014) and NBQX-treatment led to a small trend towards broader APs (Fig. 1D; P = 0.26). Importantly, sodium channel block and synaptic block both increased spontaneous AP firing frequency, indicating that homeostatic adaptations other than AP broadening were shared between both induction methods (Fig. 1E, F; P = 0.008 for control against Kyn, P = 0.005 for control against NBQX, P = 0.013 for control against TTX). These data indicate that AP broadening occurred upon sodium channel block in CA3 neurons in organotypic slice cultures and upon synaptic block in dissociated hippocampal cultures, but not in the commonly-studied model of homeostatic plasticity using dissociated hippocampal cultured neurons and 48 hours of TTX application.

### Stable AP duration in primary dissociated neurons during TTX-induced homeostatic plasticity

In contrast to our finding in dissociated cultured neurons of the hippocampus, a recent study described TTX-induced AP broadening in dissociated cultured neurons of the neocortex^13^. To resolve this discrepancy, we set out to replicate AP broadening in neocortical cultured neurons. Given that exact reproduction of experimental conditions is challenging and neuronal plasticity depends on brain region^18^, animal species^19^, and animal strain^20^, we probed homeostatic AP broadening under different experimental conditions varying: 1) experimental animal species (rat and mouse), 2) strain (C57BL/6 and CD1 mice), 3) sub-strain (C57BL/6N and C57BL/6J mice), 4) brain area (cortex and hippocampus), 5) cortical subdomain (prefrontal cortex), 6) culture duration (10–14, 15–21, and 32 days *in vitro*), 7) culture growth medium, 8) recording solutions, and 9) laboratory and experimenter performing recordings (conditions I–X in Fig. 2B, condition XI are recordings from Fig. 1; see also Fig. 2 – figure supplement 1). Even though AP durations varied up to 2-fold between conditions, statistically significant homeostatic AP broadening was not detectable in any of the tested conditions (Fig. 2B). To minimize type II errors (false negative) we intentionally did not apply a correction for multiple comparisons. The only significance was observed in condition III but in an opposite direction (i.e. AP narrowing with TTX, P=0.026; Fig. 2B). However, this is likely a false positive because application of corrections for false discovery rate results in P=0.268 for both Benjamini–Hochberg and Bonferroni correction. To detect cross-conditional AP broadening by TTX, we merged all conditions based on their difference in AP duration after TTX-treatment (Fig. 2C). However, AP duration was the same in control and TTX-treated cells (217 and 197 cells, respectively, P = 0.83).

**Figure 2.**
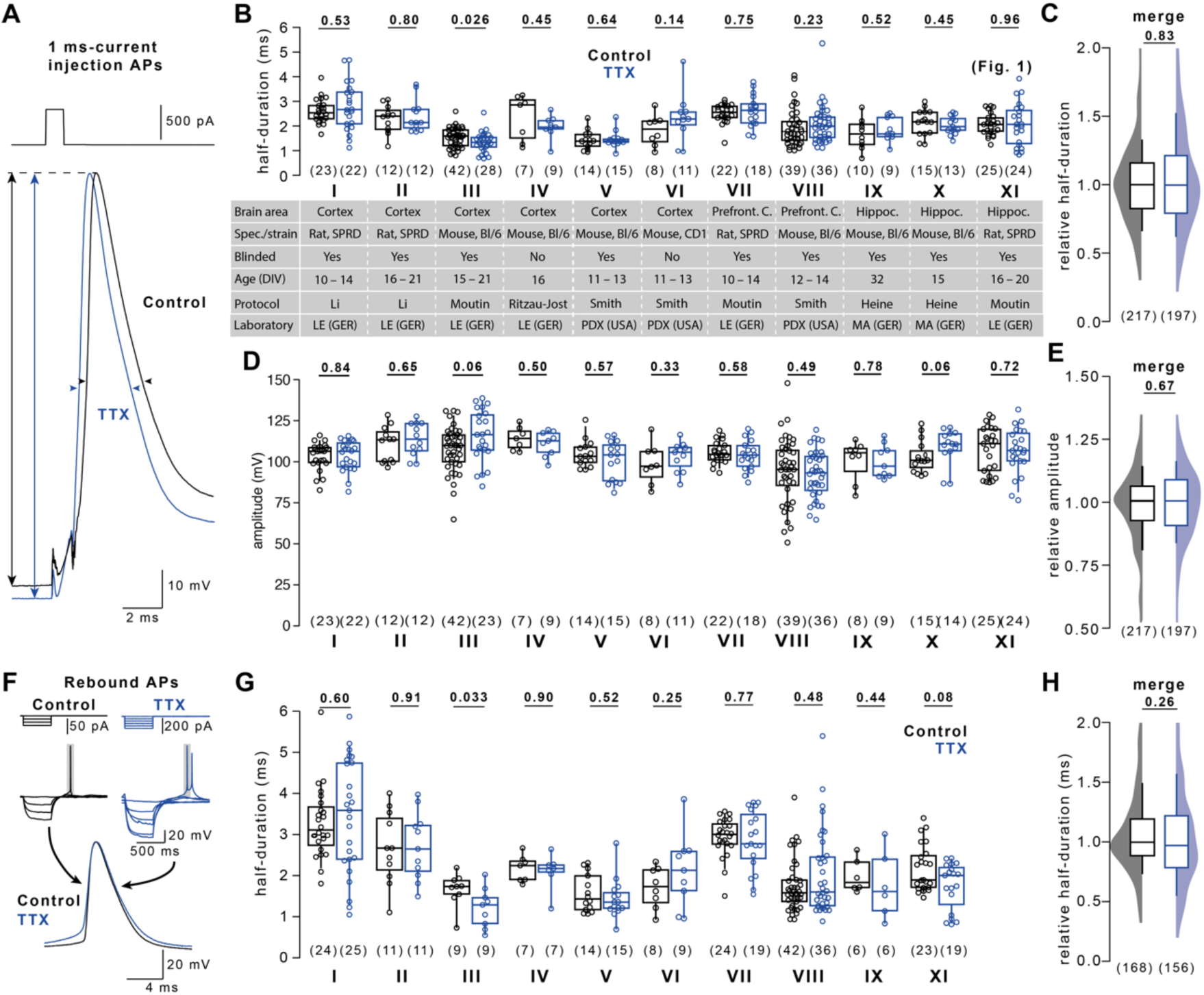
Stable AP duration in primary dissociated neurons during TTX-induced homeostatic plasticity. (A) Example current injection-evoked APs (500 pA for 1 ms) under control condition (black) and following 48 hours of TTX-treatment (blue). AP amplitudes are recorded between the membrane potential preceding the AP and AP peak (arrows), AP durations are recorded at half-maximal amplitude (arrow heads). (B) AP duration in control and TTX-treated cells (color code as in A) for eleven different experimental conditions (I–XI). Specification of individual conditions is provided in Table S1. Condition XI adopted from Fig. 1D. (C) Merge of AP durations in control and TTX-treated cells across conditions I–IX normalized to the corresponding condition’s control median duration. (D) AP amplitudes in control and TTX-treated cells as in B. (E) Cross-conditional merge of AP amplitudes in control and TTX-treated cells as in C. (F) Example hyperpolarization-evoked “rebound APs” (-150 pA or -60 pA for 500ms) in control and TTX-treated cells (color code as in A). (G) Rebound AP duration in control and TTX-treated cells as in B. (H) Cross-conditional merge of normalized durations as in C for rebound APs. Numbers in brackets reflect number of recorded neurons. Box plots as median ± interquartile range, whiskers indicate 10–90 percentile. P-values provided above graphs; significance tested using Mann-Whitney-*U* tests.

We next tested whether AP broadening was concealed by systematic differences in neuronal health or recording quality between control and treatment group. Because unhealthy neurons tend to have small and slow APs, possibly due to changes in resting membrane potential or expression of voltage-gated sodium and potassium channels, we first analyzed AP amplitude as a measure of neuronal viability. AP amplitudes were not affected following TTX treatment in any of the eleven recording conditions (Fig. 2D) or a cross-conditional comparison (Fig. 2E). In addition, we focused on the culture protocol or cortical subdomain previously associated with AP broadening^13^ (conditions I, VII, and VIII) and applied further inclusion criteria to rigorously discriminate healthy excitatory neurons: the occurrence of multiple APs upon 200 ms-current injections (≥ 2 APs), a resting membrane potential below – 55 mV, and a maximum AP depolarization rate between 100 and 337 V/s (includes 96% of excitatory and exclude >80% of the inhibitory neurons; ref. 21). Applying any or all three of these criteria did not reveal AP broadening upon TTX treatment (Fig. 2 – figure supplement 2). These data indicate that stricter exclusion criteria result in smaller variability in action potential duration but do not uncover AP broadening in these datasets.

Instead of depolarization-evoked APs, Li *et al.*^13^ recorded hyperpolarization-evoked ‘rebound APs’ (Fig. 2F), possibly reducing the impact of different resting membrane potentials and depolarizing current injection on the AP waveform. We determined that the duration of rebound APs was similarly unaffected by TTX treatment across all recorded conditions (Fig. 2G, H). These data show that TTX-induced AP broadening does not robustly occur in dissociated cultured neurons.

### Homeostatic plasticity increases network activity and excitatory synaptic transmission

To verify successful induction of homeostatic plasticity under our experimental conditions, we quantified changes in neuronal activity and synaptic strength following TTX-induced homeostatic plasticity. TTX-treatment increased the number of spontaneously active neurons (Fig. 3A; merge of conditions I–IV and VII; control: 23 out of 87 cells, TTX-treatment: 58 out of 74 cells, P < 0.001 with Fisher’s exact test) and the number of APs per spontaneous burst (Fig. 3B, C; median [IQR] 1.0 [1.0–1.12] APs and 1.67 [1.0–2.71] APs for control and for TTX-treatment, 23 and 58 cells, respectively, P < 0.001). These results indicate an adaptive increase in neuronal activity following TTX-treatment, in line with previous findings on homeostatic plasticity^16,22,23^. Furthermore, spontaneous excitatory postsynaptic currents (sEPSCs; spontaneous synaptic currents recorded in the absence of TTX) were recorded as a measure of synaptic strength (Fig. 3D). sEPSC frequency and amplitude increased upon TTX-treatment (Fig. 2E, merge of recordings from condition I and II; median [IQR] frequency 1.11 [0.42–1.47] Hz and 5.07 [1.95–5.52] Hz in control and TTX-treated cells, 16 and 11, respectively, P = 0.027; median [IQR] amplitude 12.4 [10.6–14.4] pA and 15.5 [14.1–17.4] pA in control and TTX-treated cells, 16 and 11 cells, respectively, P = 0.042), in accord with known homeostatic synaptic adaptations to inactivity^24–27^. These changes in sEPSC amplitude and frequency are not specific for somatic, pre-or postsynaptic adaptations. However, the results show that blocking AP firing with TTX successfully induced homeostatic plasticity under our experimental conditions.

**Figure 3.**
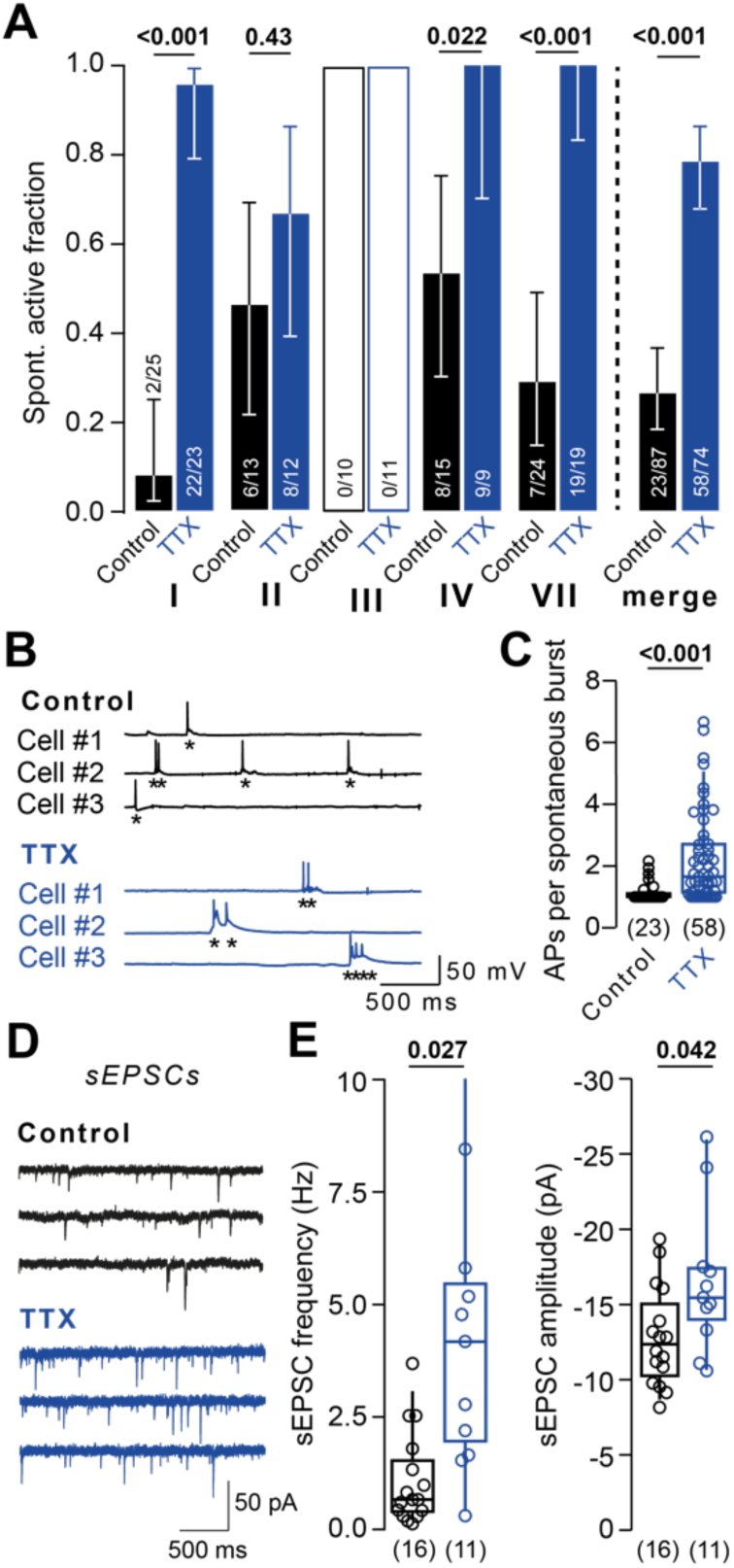
Homeostatic plasticity increases network activity and excitatory synaptic transmission. (A) *Left:* Spontaneously active control (black) and TTX-treated cells (blue) in condition I–IV and VII. *Right*: Active neurons merged across conditions I–IV and VII. (B) Example spontaneous AP bursts in control and TTX-treated cells. Asterisks mark individual APs. (C) AP number per burst in control and TTX-treated cells across conditions I–IV and VII. (D) Example spontaneous excitatory postsynaptic currents (sEPSCs) in control and TTX-treated cells. (E) *Left:* sEPSC frequency and *Right:* sEPSC amplitude in control recordings and after TTX-treatment (merged data from conditions I and II). Numbers in brackets reflect number of recorded neurons. Bars in A reflect fraction of spontaneously active neurons. Whiskers in A reflect confidence limits for 95% confidence interval based on Wilson interval. Box plots as median ± interquartile range, whiskers indicate 10–90 percentile. P-values above graphs, significance tested by Fisher’s exact test in A or Mann-Whitney-*U* test elsewhere.

### Lack of BK channel-dependent AP broadening in dissociated cultured neurons

Various potassium channel subtypes regulate AP repolarization and could therefore underlie AP broadening. Li *et al.*^13^ reported that downregulation of BK-type potassium channels led to homeostatic AP broadening in neocortical cultured neurons. We tested the role of BK channels in AP repolarization (neurons grown under condition I). However, blocking BK channels with Iberiotoxin (IbTx, 300 nM) did not lead to broadening of depolarization-evoked APs in control and TTX-treated cells (Fig. 4A; P = 0.10 and 0.48, respectively), or a merge of both conditions (Fig. 4B; P = 0.10). Rebound APs were similarly unaffected by BK channel block (Fig. 4 – figure supplement 1). Next, we tested whether recurrent neuronal activity, which increases cytosolic calcium and boosts BK-channel activity, enhanced AP broadening following TTX treatment. While APs significantly broadened during repetitive firing, AP durations were similar in control and TTX-treated neurons (Fig. 4C and D), suggesting no differential regulation of BK-channels following TTX-treatment in these recordings.

**Figure 4.**
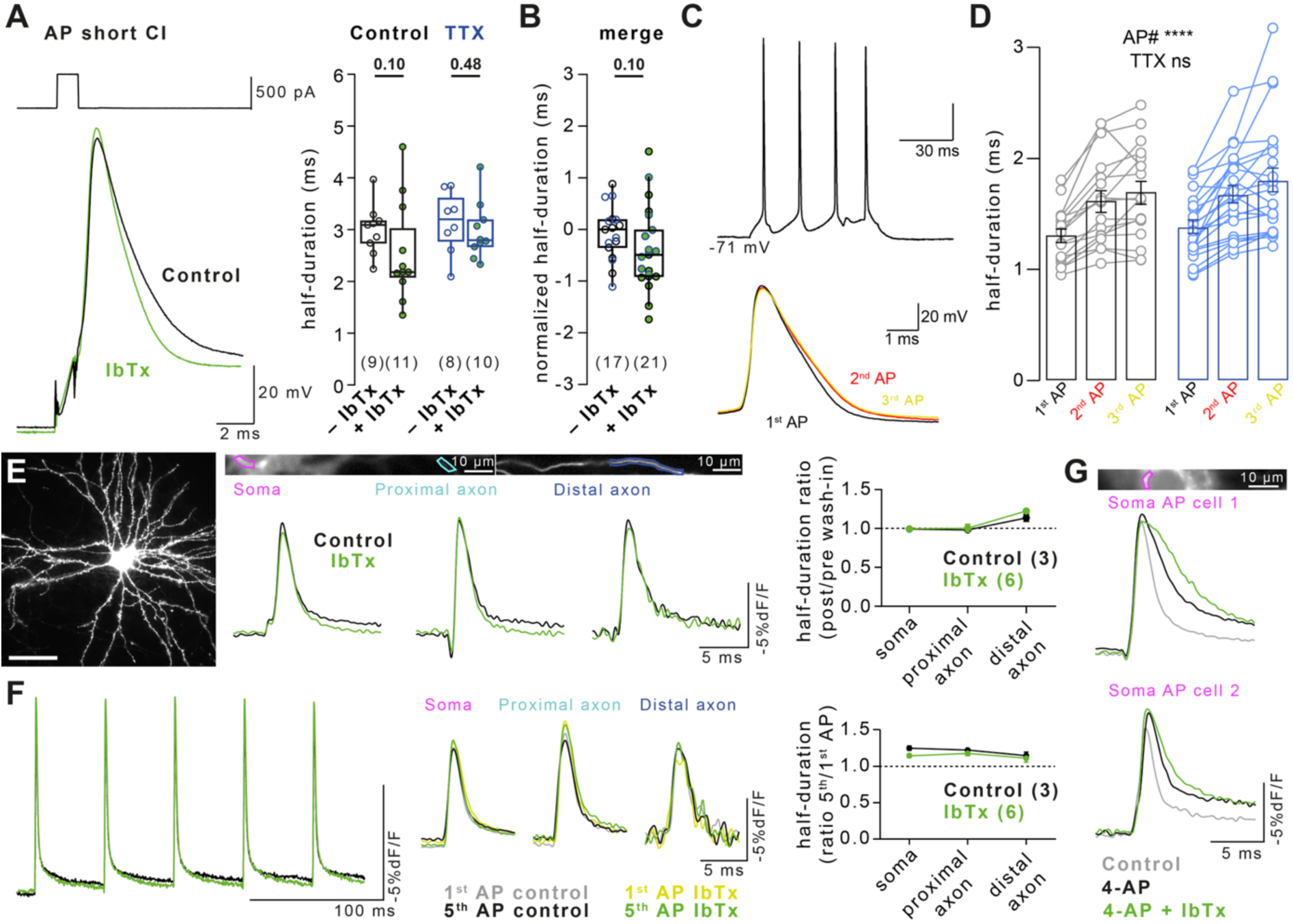
Lack of BK channel-dependent AP broadening in dissociated cultured neurons. (A) *Left:* Example current injection-evoked APs (500 pA for 1 ms) in control solution (black) and solution containing 300 nM Iberiotoxin (IbTx, green). *Right:* AP duration in control and TTX-treated cells recorded IbTx-free (black and blue, respectively) or IbTx-containing solution (green). Data recorded under condition I. (B) Merged AP duration across control and TTX-treated cells for recordings in IbTx-free and IbTx-containing solution (data in A normalized to the respective group’s median duration in control solution). (C) APs evoked by a 200 ms current injection (top) and overlay of the forst three evoked APs (bottom).(D) Broadening of AP duration of the 3 first APs during 200 ms current injections in control (black, *left*) and TTX-treated cells (blue, *right*). Bars as mean ± SEM, lines with markers depict recordings from individual neurons. (E) *Left:* Example image of neuron expressing the genetically encoded voltage indicator Ace-mNeon (scale bar 50 µm). *Middle:* Example Ace-mNeon-recorded somatic, proximal axonal, and distal axonal APs in a primary hippocampal cultured neuron in IbTx-free solution (black) and following BK channel block by IbTx (100 nM, green). *Right:* Ratio of somatic and axonal AP durations in IbTx-free and IbTx-containing solution. (F) *Left:* Example AP train (5 x 20 Hz) and overlay of 1^st^ and 5^th^ train AP at the soma, proximal, and distal axon in control condition (grey and black) and following BK channel block by IbTx (yellow and green). *Right:* AP broadening (ratio of 5^th^ over 1^st^ train AP duration) in the three compartments in control and IbTx-containing solution. (G) Somatic APs under control condition (grey), following treatment with 4-AP (black), and following treatment with 30µM 4-AP + 100 nM IbTx (green). Numbers in brackets reflect number of recorded neurons. Box plots as median ± interquartile range, whiskers indicate 10–90 percentile. Plots in D–F as mean ± SEM. P-values above graphs, significance tested by Mann-Whitney-*U* test.

In electrical recordings, intracellular recording solution may have disturbed cytosolic calcium dynamics and therefore BK- and thus calcium-dependent AP repolarization. As an independent approach, we recorded the AP duration in hippocampal cultured neurons using the genetically encoded voltage indicator (GEVI) Ace-mNeon^28^ before and after acute IbTx perfusion (100 nM). BK channel block did not affect AP duration in the soma or along the axon (Fig. 4E). Using the GEVI, we again tested the role of BK channels in AP broadening during a train of APs (5 APs at 20 Hz, Fig. 4F). The broadening of somatic, proximal axonal, and distal axonal APs during trains was similar before and after BK channel block, indicating that AP duration was not modulated by BK channels. To confirm that IbTx was indeed pharmacologically active in Ace-mNeon recordings, we preconditioned cells with 4-AP (30 µM) prior to IbTx perfusion (Fig. 4G). Neurons preconditioned with 4-AP had broader somatic APs which were further broadened by IbTx, presumably due to increased calcium influx following 4-AP treatment, in turn activating more BK channels^29^. These data indicate that IbTx application was pharmacologically active but BK channels did not contribute to AP repolarization in cultured neocortical neurons.

## Discussion

Homeostatic AP broadening has been reported in some systems, but it has not been tested systematically across experimental models so far. Because BK channels that were proposed to underlie AP broadening are activated by depolarization and calcium^30,32^, we studied APs elicited with depolarizations from the RMP where BK activity was presumably increased, as well as anode break (‘rebound’) APs thought to obviate variance from ion channel inactivation. We found that broadening of rebound and depolarization-evoked APs did not occur following the induction of homeostatic plasticity with TTX in the most commonly studied model system (primary dissociated neuronal cultures). However, AP broadening was observed in cultured hippocampal slices. Furthermore, blocking BK channels, which were reported to mediate AP broadening during homeostatic plasticity^13^, did not impact AP duration in dissociated cultured neurons. Our results therefore indicate that AP broadening in cultured dissociated neurons is not a required component of homeostatic plasticity.

### TTX-induced homeostatic AP regulation in organotypic hippocampal cultured neurons

AP broadening was previously shown to occur in CA3 neurons of organotypic hippocampal slice cultures following TTX-induced homeostatic plasticity^16^. Consistently, we recorded broader APs in CA3 neurons following neuronal inactivity (Fig. 1). In contrast to CA3 neurons, granule cells of the dentate gyrus did not show TTX-induced AP broadening^17^. Similarly, CA1 neurons did not show AP broadening following synaptic block-induced homeostatic plasticity^31^. Therefore, we conclude that homeostatic AP broadening in organotypic hippocampal cultures is most likely cell type-specific.

### TTX-induced homeostatic AP regulation in primary dissociated cultured neurons

To the best of our knowledge, only three studies so far reported on the duration of APs following TTX-induced homeostatic plasticity in dissociated cultured neurons^13,33,34^. While Li *et al.*^13^ observed TTX-induced broadening, the studies by Lee, Chung, and others^33,34^ reported stable AP duration, consistent with our findings (Fig. 2). Furthermore, in other studies investigating TTX-induced homeostatic plasticity in dissociated cultured neurons, prominent AP broadening was either missed or absent (see for example refs. 23,35–37). One possible explanation for this inconsistency could be the cell type-specificity of AP broadening. We focused on large, presumably excitatory neurons. However, we and others could have selected different neuron types compared to Li *et al.*^13^. In an attempt to reduce bias, we incorporated multi-lab, randomization, and blinding strategies across many of our experiments. Only recordings with leaky seal formation or inadequate capacitance compensation^38^ were excluded. However, in a subset of experimental conditions we applied additional exclusion criteria to ensure that variability in neuron type, health or recording quality were not causing a type II (false negative) statistical error. Application of these exclusion criteria did not reveal AP broadening in our data (Fig. 2 – figure supplement 2), confirming that AP broadening was not associated with homeostatic plasticity in neocortical neurons following TTX treatment.

The sensitivity to chronic TTX treatment might depend on baseline neuronal activity, which is in part related to neuronal culture density^39^. However, TTX did not induce AP broadening despite different baseline activities (Fig. 3A) and a nearly threefold variation in the number of plated cells per cover slip between conditions (25k – 70k cells per cover slip). Although we mainly focus on neocortical cultured neurons (condition I to VIII, Fig. 2) because Li *et al.* used neocortical neurons, the absence of AP broadening in hippocampal neurons (group IX to XI) could in principle be explained by the selective loss of CA3 neurons, which show AP broadening in organotypic cultured neurons (Fig. 1A and B). However, CA3 neurons were shown to survive in dissociated cultures following region-specific microdissection^40^, and CA1 neurons are generally more stress-sensitive to excitotoxicity with glutamate or NMDA than CA3 and DG neurons^42^, arguing against a general selective loss of CA3 neuron in dissociated cultures. Altered cell-cell interactions with glia and neurons in organotypic and dissociated neuronal cultures could instead contribute to the different findings in various hippocampal preparations.

### Synaptic block-induced homeostatic AP regulation in primary dissociated cultured neurons

Homeostatic AP broadening was shown in CA3 neurons upon activity block by the low-affinity glutamate receptor blocker Kyn^15^. We similarly observed Kyn-induced AP broadening in dissociated cultured hippocampal neurons (Fig. 1C, D). In contrast, APs were not significantly broader following synaptic block by NBQX (Fig. 1C, D), in accord with recordings from CA1 neurons in organotypic cultures using CNQX. TTX-induced broadening may therefore be cell-type specific or due to a differential effect of the glutamate receptor blockers on NMDA receptors which are blocked by Kyn but not NBQX/CNQX or TTX and which have recently been demonstrated to be important for the induction of synaptic homeostatic plasticity^41^. Further studies are needed to investigate the role of subcellular calcium signals in the regulation of AP duration in hippocampal neurons during block of glutamate receptors.

### Absence of BK channel-dependent AP broadening in dissociated cultured neurons

Li *et al.*^13^ proposed that BK channels hasten AP repolarization and that this effect is lost during TTX-induced homeostatic plasticity, leading to AP broadening. We did not detect AP broadening when blocking BK channels in either control or TTX-treated cells (Fig. 4). Furthermore, because calcium accumulation following multiple APs has been shown to enhance intraspike BK contribution in some neurons^43^ we compared how 1^st^, 2^nd^, or 3^rd^ APs were impacted by prior TTX-exposure. AP broadening did occur in the later APs but the changes were unaffected by TTX exposure (Fig. 4D), suggesting no activity-dependent effect of TTX or BK channels on AP duration.

Voltage imaging using Ace-mNeon similarly showed that duration of single APs and broadening during AP trains in the soma and axon were unaffected by BK channel block. Blocking BK channels only affected AP repolarization when cells were preconditioned with 4-AP. These results are consistent with previous reports showing that in some neurons BK channels do not contribute to repolarization^43,44^ while doing so in others^44–47^. Differences in the abundance of other types of potassium channels^44^ and intracellular calcium buffering^46^ are known to affect the contribution of BK channels to AP repolarization and may account for the lack of AP broadening upon BK channel block in our dissociated cultured neurons. Alternatively, BK activity may occur only under conditions associated with elevated intracellular^30,32^. The lack of BK channel contribution to AP repolarization in our dissociated neurons is another possible explanation (in addition to different selection of cells) for why we and others have not observed the homeostatic AP broadening described by Li *et al.*^13^.

### Inactivity-induced homeostatic AP regulation *in vivo*

Homeostatic plasticity has also been studied using *in vivo* recordings from the activity-suppressed visual cortex^48,49^. The majority of these studies reported stable AP duration upon monocular deprivation or chemogenetic activity suppression^50–52^. In contrast, Li *et al.*^13^ found homeostatically broadened APs in the visual cortex upon monocular deprivation. Furthermore, stable AP duration was also observed in other brain areas following activity suppression such as (1.) the somatosensory cortex upon unilateral whisker trimming^53–58^ or forepaw denervation^59^, (2.) the auditory system upon cochlear ablation^60,61^, hearing loss^62^, or congenital deafness^63^, (3.) the vestibular nucleus upon vestibular ablation^64^, and (4.) the olfactory mitral cells upon naris occlusion^65^ or senescent hippocampal deafferentiation^66^. Besides somatic APs, APs in excitatory presynaptic terminals of the somatosensory thalamus were also unaffected by whisker deprivation^67^. Thus, AP broadening does not appear to occur robustly upon suppression of neuronal activity in various neuronal pathways.

### Hyperactivity-induced homeostatic AP regulation

Homeostatic plasticity occurs not only upon activity block but also upon hyperactivity, generally involving opposite adaptations. In cultured neurons, AP duration was unaffected following hyperactivity induced by chronic depolarization^68^, block of potassium channels^69^, block of inhibitory synaptic transmission^31,33,34^, or chronic ChR2-induced photostimulation^70^. In contrast, Li *et al.*^13^ reported similar BK channel downregulation upon both, TTX-induced activity block and depolarization-induced hyperactivity.

As in cultured neurons, AP duration was mostly unaffected in brain slice recordings from animals with increased neuronal activity due to an enriched environment^58,71,72^, psychosocial stress^73^, or febrile seizures^74^. AP shortening occurred in cerebellar granule cells after exposure to an enriched environment^75^, in auditory neurons following chronic noise exposure^76^, and in hippocampal neurons following age-related^77^ or pilocarpine-induced hyperactivity^78^. Thus, in contrast to Li *et al.*^13^, AP duration was either unaffected or, if anything, shortened during hyperactivity-induced homeostatic plasticity in previous studies.

In this study, several laboratories collaborated to determine the robustness of inactivity-induced AP broadening. AP broadening did not occur upon TTX-treatment in dissociated hippocampal or neocortical neurons. Furthermore, BK channel block did not affect AP duration in dissociated neurons, arguing against BK channels as mediators of a general homeostatic AP regulation.

Despite the lack of homeostatic, TTX-induced AP broadening in dissociated cultures, AP duration was broadened upon Kyn-treatment in dissociated cultures and using TTX in CA3 neurons in organotypic cultures. Because BK-channels control AP duration in CA3 neurons of organotypic cultures^79^, homeostatic BK-channel downregulation as proposed by Li *et al.* may be involved in AP broadening in this specific cell type. While the reasons for the variable occurrence of homeostatic AP broadening remain unknown, this may render neuronal circuitries more robust to perturbations. The regulation of AP duration therefore might represent one element in the repertoire of neuronal plasticity that is, similar to other plasticity mechanisms, not generally shared, but specifically expressed in some cell types and neuronal compartments.

## METHODS

### Ethics

At all times animal procedures were in accordance with the relevant international, national and institutional guidelines for the care and use of laboratory animals (European Council Directive 86/609/EEC, U.S. Public Health Service Policy on Humane Care and Use of Laboratory Animals, U.S. National Institutes of Health Guide for the Care and Use of Laboratory Animals, German Tierschutzgesetz, French National Research Council guidelines, and Leipzig and Mainz University guidelines). All animal procedures were approved in advance by the local health authorities (Ethics Committees at Leipzig University and Mainz University, federal Saxonian Animal Welfare Committee, the District administration Mainz-Bingen (41a/177-5865-§11 ZVTE), the Committee of the Préfecture des Bouches-du-Rhône (D13055-08), V.A. Portland Health Care System Institutional Animal Care and Use Committee).

### Animal models

Organotypic hippocampal cultures were generated from postnatal day 5–7 Wistar rats. Dissociated neocortical cultures were generated from postnatal day 0 or 1 Sprague-Dawley rats (conditions I, II and VII), C57BL/6N mice (conditions III and IV), C57BL/6J mice (conditions V and VIII), or CD1 mice (condition VI). Dissociated hippocampal cultures were generated from postnatal day 0 or 1 C57BL/6N mice (conditions IX), C57BL/6J mice (condition X), or Sprague-Dawley rats (condition XI). Generally, animals of both sexes were used to generate cultures. Mother animals were kept in individually ventilated cages under a 12h/12h light-dark cycle and received water and food *ad libitum*.

### Induction of Homeostatic Plasticity

Homeostatic plasticity was induced by inclusion of one of the respective blockers in culture medium for 48–56 hours prior to recordings. All blockers were first diluted in 100 µL of fresh medium per treated well and subsequently added to the respective wells containing 1 mL of medium. For control conditions, only 100 µL of fresh medium without blockers was added to the respective wells. The blockers had final concentrations of (in mM): 0.002 tetrodotoxin (TTX; Tocris, Wiesbaden-Nordenstadt, Germany), 2 kynurenic acid (Kyn), or 0.01 2,3-Dioxo-6-nitro-1,2,3,4-tetrahydrobenzo-[f]quinoxaline-7-sulfonamide (NBQX). To avoid bias, coverslips were randomized to treatment (control or blocker) and recordings were made and analyzed in a blinded fashion (except for condition IV and VI). Blinding was performed by an independent investigator. Control or TTX-treatment groups were assigned using an 8-block randomization approach for experiments of condition VIII. Unblinding of the experimental groups was performed after analysis and disclosure of the data by the experimenter.

### Preparation of dissociated neocortical cultures

#### Conditions I and II (Hallermann Lab, Leipzig, Germany)

Neocortical cultures were generated as described by Li *et al.*^13^. Shortly, neocortices were isolated and dissected into 5–10 pieces per hemisphere. Samples were washed twice in ice-cold modified HBSS (4.2 mM NaHCO_3_ and 1 mM HEPES, pH 7.35, 300 mOsm) supplemented with 20% fetal bovine serum (FBS) and digested at 37 °C for 30 min using papain. After stopping digestion with 20% FBS-containing modified HBSS, samples were washed and dissociated using Pasteur pipettes, pelleted twice, and filtered using a 70 nm nylon strainer. 30.000–50.000 vital dissociated cells were plated on poly-D-lysine-coated 14 mm coverslips and cultured in NbActive4 medium (Transnetyx, Cordova, TN, USA) in an incubator until use (37 °C, 5% CO_2_ in room air, water vapor saturated).

#### Conditions III and VII (Hallermann Lab, Leipzig, Germany)

Cultures were generated as previously described by Moutin *et al.*^80^. In brief, mice were decapitated and neocortices (for condition III) or the frontal third of neocortices (for condition VII) were isolated in ice-cold Hibernate-A medium (Thermo Fisher Scientific, Waltham, MA, USA) and cut into 5–10 chunks. Chunks were first digested by papain (dissolved in Hibernate-A; 10 min at 37 °C) and subsequently by papain + DNAse (125 µL of 20 mg/mL DNAse stock; 5 min at 37 °C). Digestion was stopped by CM+ medium (Neurobasal A 86.5 %, B27 2 %, Glutamax 0.25 %, L-Glutamine 0.25%, penicillin G/streptomycin 1 %, heat-inactivated FBS 10 %) and chunks were mechanically dissociated in two steps by 1250 µL pipette tips with gravity decantation and collection of the supernatant between dissociation steps and after dissociation. Supernatant was passed through 2 mL of frothed bovine serum albumin, centrifuged (7 min, 300 x g), the resulting pellet was resuspended in 1–2 mL of CM+ medium and 25.000–30.000 resuspended vital cells were plated onto poly-L-lysine-coated coverslips. After 30 min, 0.5 mL of CM+ medium was added to the wells and after 2 days *in vitro* (DIV), cytosine arabinoside (ARA-C, 1 µM final concentration) was added to the wells for 8–18 h. Subsequently, 0.4 mL medium per well was replaced by fresh CM- medium (BrainPhys 96.75% by Stemcell Technologies, Vancouver, Canada, B27 2%, Glutamax 0.25%, penicillin G/streptomycin 1%) and further 0.1 mL of medium were exchanged every 6–7 days with fresh CM- medium until cells were used.

#### Condition IV (Hallermann Lab, Leipzig, Germany)

Cultures were generated following the protocol by Ritzau-Jost *et al.*^14^. In short, mice were decapitated and cerebral cortices were removed, dissected in ice-cold Hank’s balanced saline solution, and cut into 5–10 pieces per hemisphere. Samples were digested by Trypsin (5 mg, dissolved in digestion solution; for composition see below) for 5 minutes at 37 °C. Trypsin was stopped by ice-cold MEM growth medium (composition see below), supernatant was discarded, and samples were mechanically and enzymatically dissociated using Pasteur pipettes of narrowing tip diameters in DNAse-supplemented MEM growth medium (10 mg/mL). The supernatant was centrifuged and the resulting pellet was resuspended in MEM growth medium for plating onto Matrigel-coated 14 mm coverslips (50.000 vital cells/coverslip). Forty-eight hours after plating, the medium was fully replaced by MEM growth medium containing 4 µM ARA-C to limit glial growth. After another 48 hours, the medium was fully replaced by MEM growth medium. Cells were incubated at 37 °C, 93% humidity, and room air plus 5% CO_2_ until use. Digestion solution contained the following (in mM): 137 NaCl, 5 KCl, 7 Na_2_HPO_4_, 25 HEPES, pH adjusted to 7.2 by NaOH. MEM growth medium was prepared from 1l MEM (Earle’s salts + L-Glutamine) supplemented with 5 g Glucose, 0.2 g NaHCO_3_, 0.1 g bovine holo-Transferrin, 0.025 g bovine insulin, 50 mL FBS and 10 mL B-27.

#### Conditions V, VI and VIII (Smith Lab, Portland, OR, USA)

Cultures were generated following the protocol by Martiszus *et al.*^81^. Briefly, 1–2-day old mice were decapitated following general anesthesia with isoflurane, and their cerebral cortices (for conditions V and VI) or the frontal third of neocortices (for condition VIII) were dissected out while bathed in ice-cold modified Hank’s balanced saline solution (modified HBSS; Hyclone laboratories, Logan, UT, USA) and cut into 5–10 pieces per hemisphere. Samples were digested by Trypsin (10 mg, dissolved in digestion solution as mentioned in Condition IV) for 5 minutes at 37 °C. Trypsin was de-activated by ice-cold HBSS with 20% FBS, supernatant was discarded, and samples were enzymatically and mechanically dissociated using Pasteur pipettes with narrowing tip diameters in DNAse-supplemented HBSS. The supernatant was centrifuged and the resulting pellet was resuspended in 5% MEM growth medium (see composition above in Condition IV) for plating 65.000 vital cells onto Matrigel-coated 12 mm glass coverslips. MEM growth medium containing 4 µM ARA-C was added 48 hours later to limit glial proliferation, and subsequently replaced with MEM growth medium following another 48 hours. Cells were incubated at 37 °C, 93% humidity, and room air plus 5% CO_2_ until use between days 11–14 *in vitro*.

### Preparation of dissociated hippocampal cultures

#### Condition IX and X (Heine Lab, Mainz, Germany)

In short, mice pups were decapitated and brains were removed. Hippocampi were dissected in ice-cold Hank’s balanced saline solution and digested by trypsin for 15 minutes at 37 °C. Next, the tissue was washed with cold HBSS and mechanically dissociated using Pasteur pipettes in DNAse-, glutamate- and FBS-supplemented DMEM growth medium. 100 µL of cell suspension containing approximately 70.000 cells was placed on a glass coverslip and incubated at 37 °C, 5% CO_2_ for 1 hour. Coverslips were then transferred to 12-well plates with glutamate-, B27- and sodium pyruvate-supplemented Neurobasal A medium (1 mL per well) and cultivated until use. Double distilled water was added in between the wells of the 12-well plate to prevent evaporation of the medium during cultivation.

#### Condition XI (Hallermann Lab, Leipzig, Germany)

Cultures were generated as described for Condition III, except that hippocampi from postnatal day 0–1 Sprague-Dawley rats were used for preparation. 25.000–30.000 vital cells were plated per coverslip.

### Recordings from dissociated cultures

Within conditions, recordings were performed blinded (except conditions IV and VI) and interleaved from randomized sister coverslips of the same culture, at the same recording setup, and by the same experimenter.

#### Conditions I – IV, VII and XI (Hallermann Lab, Leipzig, Germany)

Voltage- and current-clamp recordings were performed with borosilicate pipettes using a HEKA EPC10/2 amplifier (HEKA Elektronik, Lambrecht/Pfalz, Germany). Borosilicate glass tubes (Science Products, Hofheim, Germany) were pulled by a DMZ Universal Electrode Puller with filament heating (Zeitz Instruments, Martinsried, Germany) to 3–6.5 MΩ pipette resistance. Pipettes were filled with internal solution (see *Solutions and reagents* section) and mounted on a micromanipulator (Kleindiek Nanotechnik, Reutlingen, Germany) using a custom-built pipette holder. Recordings were performed at room temperature (22–25 °C). All current-clamp recordings were filtered with the internal 10 kHz 8-pole Bessel filter of the HEKA EPC10/2 amplifier and subsequently digitized (200 kHz) with the HEKA EPC10/2 using Patchmaster software (HEKA Elektronik, Lambrecht/Pfalz, Germany). Built-in compensation was used for voltage offset and pipette capacitance. Membrane potentials were not corrected for liquid junction potential and bridge compensation was adjusted to minimize the current injection artifact. APs were evoked by brief 1 ms depolarizing current injections of 500–1500 pA or 500 ms hyperpolarizing pulses (300–1000 pA) leading to APs upon return to resting membrane potential (‘rebound APs’). Both, depolarization-evoked and rebound APs were first recorded at resting membrane potential without holding current. While depolarization readily evoked APs in all conditions, rebound AP did not occur in all conditions at their resting membrane potential. In these cells, holding currents were used to depolarize cells until rebound APs were observed upon long hyperpolarizing pulses. Spontaneous excitatory postsynaptic currents were recorded in voltage-clamp mode at -70 mV membrane potential in neurons grown in conditions I and II. Currents were filtered with the internal 3 kHz 8-pole Bessel filter of the HEKA EPC10/2 amplifier, digitized (200 kHz) and detected using the template matching algorithm ^82^ implemented in NeuroMatic (Version 2.00, 15.09.2008; ref. ^83^).

#### Conditions V, VI and VIII (Smith Lab, Portland, OR, USA)

Current-clamp recordings were performed with borosilicate pipettes using a HEKA EPC10 amplifier (HEKA Elektronik, Lambrecht/Pfalz, Germany). Patch pipettes (5–10 MΩ resistance) were pulled from thin-walled borosilicate glass tubes (Warner Instruments, Holliston, MA, USA) using a P-87 electrode puller (Sutter Instruments, Novato, CA, USA). Pipettes were filled with an internal solution (see *Solutions and reagents* section) and mounted on a micromanipulator (ROE-200, Sutter Instruments, Novato, CA, USA). Recordings were performed at room temperature (22–25 °C). Voltage signals were filtered at 2.9 kHz and digitized at 20 kHz. Membrane potentials were corrected for liquid junction potential and bridge balance was performed manually prior to data acquisition. All APs were evoked by a sequence of transient injections relative to the resting membrane potential and measurements taken from the APs elicited with the minimal stimulus using either depolarizing current injections (500–1000 pA in 100 pA increments for 1–3 ms in 0.5 ms increments in conditions V and VI or 20–140 pA in 20 pA increments for 0.5 s in condition VIII) or following rebound from longer hyperpolarizing pulses (300–1200 pA in 100 pA increments for 500 ms).

#### Conditions IX & X (Heine Lab, Mainz, Germany)

Voltage- and current-clamp recordings were performed with borosilicate pipettes using a HEKA EPC9 amplifier (HEKA Elektronik, Lambrecht/Pfalz, Germany). Filamented glass tubes (Science Products, Hofheim, Germany) were pulled by a Narishige PC-100 electrode puller. Resistance of the pipettes was 2–6 MΩ. Pipettes were filled with internal solution (see *Solutions and reagents* section) and mounted on a micromanipulator (Luigs & Neumann, Ratingen, Germany). Recordings were performed at room temperature (21–24 °C). Signals were filtered at 2.9 kHz and digitized at 20 kHz using Patchmaster software (HEKA Elektronik, Lambrecht/Pfalz, Germany). Built-in compensation was used for voltage offset and pipette capacitance. Membrane potentials were not corrected for liquid junction potential. APs were evoked by brief 1 ms depolarizing current injection of 50–300 pA or 500 ms hyperpolarizing pulses (300–400 pA) leading to APs upon return to resting membrane potential (‘rebound APs’).

### Preparation of organotypic hippocampal cultures

Cultures were prepared as described previously^84^. In brief, young Wistar rats (P5–P7) were anesthetized with isoflurane and killed by decapitation, the brain was removed, and each hippocampus was dissected. Hippocampal slices (350 μm) were obtained using a Vibratome (Leica, VT1200S, Wetzlar, Germany). They were placed on 20 mm latex membranes (Millicell; Merck, Darmstadt, Germany) inserted into 35 mm Petri dishes containing 1 mL of culture medium and maintained for up to 8 days in an incubator at 34 °C, 95% O_2_, 5% CO_2_. The culture medium contained 25 mL MEM, 12.5 mL HBSS, 12.5 mL horse serum, 0.5 mL penicillin/streptomycin, 0.8 mL glucose (1 M), 0.1 mL ascorbic acid (1 mg/mL), 0.4 mL HEPES (1 M), 0.5 mL B27, and 8.95 mL sterile H_2_O. To limit glial proliferation, 5 μM ARA-C was added to the culture medium starting at DIV 4.

### AP recordings from organotypic hippocampal cultures

Whole-cell recordings were obtained from CA3 neurons in organotypic cultures at DIV 7 to DIV 10. All recordings were made at 34 °C in a temperature-controlled recording chamber (Luigs & Neumann, Ratingen, Germany) using 7–10 MΩ patch pipettes. CA3 pyramidal neurons were recorded in current-clamp with a Multiclamp 700B Amplifier (Molecular Devices, San Jose, CA, USA). APs were elicited by delivering DC step pulses (duration 1.0 s) at an interstimulus interval of 10–15 s. During the whole recording, ionotropic glutamate and GABA_A_ receptors were blocked with 2–4 mM kynurenic acid and 100 μM picrotoxin, respectively.

### Analysis of APs

Data were analyzed using IGOR PRO software (WaveMetrics, Lake Oswego, OR, USA; Version 6.32A or 6.37), the Patcher’s Power Tool (Version 2.19 Carbon, 15.03.2011) and NeuroMatic (Version 2.00, 15.09.2008; ref. ^83^) extensions, as well as self-written IGOR PRO scripts. AP durations were quantified as the duration at half-maximal AP amplitude, amplitude being the difference between AP peak and the membrane potential preceding stimulation. When AP durations were recorded at half-maximal amplitude between AP threshold (defined as the membrane potential at which the voltage rate of change first reaches 10 V/s) and AP peak, AP durations were still unaffected by TTX treatment (data not shown). To exclude the effect of mis-balanced bridge compensation on AP shape, only APs that occurred after the end of the second current injection artifact (elicited by the end of current-injection) were analyzed. In some recordings, current injection artifacts were blanked over 50 µs at the beginning and the end of the current injection and only experiments in which blanking did not affect AP amplitude and duration were subsequently analyzed.

### Hippocampal dissociated cultures and transfection for GEVI imaging

Culture preparation and GEVI transfection were performed as previously described by Sabater *et al.*^28^. In short, hippocampi were dissected from embryonic day 17.5 Wistar rat pups of either sex, treated with trypsin (Worthington Biochemical Corporation, Lakewood, NJ, USA) at 0.5 mg/mL, and mechanically dissociated using fire polished Pasteur pipettes. Neurons were plated on 18 mm glass coverslips pre-treated with 100 µg/mL poly-L-lysine and coated with 10 µg/mL laminin (Thermo Fisher Scientific, Waltham, MA, USA). Cultures were maintained in Neurobasal medium with B27 (1x) and GlutaMAX (1x, all obtained from Thermo Fisher Scientific, Waltham, MA, USA), supplemented with FBS (2%, Biosera, Nuaille, France) and Penicillin/Streptomycin (1%, Sigma-Aldrich, St. Louis, MO, USA), at 37°C in a humidified incubator with 5% CO_2_. For transfections with Ace2N-mNeon-4AA (Ace-mNeon, ref. ^85^; distributed by Biolife Solutions Inc., Bothell, WA, USA), the Effectene transfection reagent (Qiagen, Venlo, Netherlands) was used. The medium was changed at DIV 3 to culture medium without antibiotic or FBS, and transfections with Effectene were performed at DIV 7 following the manufacturer’s protocol. After transfection neurons were maintained in serum-free media without antibiotics. For all experiments 30% of the medium was changed weekly and neurons were imaged 7–14 days after transfection (14–21 DIV).

### GEVI Imaging conditions

GEVI imaging was performed as previously described by Sabater *et al.*^28^. In short, neurons were imaged using an inverted Olympus IX71 epifluorescence microscope with a 60x 1.42 NA oil-immersion objective (Olympus, Tokyo, Japan). Coverslips were mounted in a heated chamber (Warner instruments, Holliston, MA, USA; total volume ∼500 µL) and placed on an IMTP microscope stage (Scientifica, Uckfield, United Kingdom). Cells were maintained in external HEPES-buffered saline solution (HBS: 2 mM CaCl_2_, 1.6 mM MgCl_2_, 1.45 mM NaCl, 2.5 mM KCl, 10 mM Glucose, 10 mM HEPES, pH 7.4, Osmolarity 290 mOsm) and experiments were carried out at near physiological temperatures of 32–35 °C.

In Ace-mNeon imaging experiments, ∼505 nm excitation illumination was provided using a 525 nm LED (Solis LED, Thorlabs, Newton, NJ, USA), a 500/20 nm excitation filter, and a 510 nm long-pass dichroic mirror. Ace-mNeon emission transmitted through the dichroic was filtered using a 520 nm long-pass filters (Chroma, Bellows Falls, VT, USA). A power density of 10 mW/mm^2^ was obtained at the specimen plane. Images were acquired at 3.2 kHz with an ORCA-Flash4.0 V2 C11440-22CU scientific CMOS camera (Hamamatsu Photonics, Hamamatsu, Japan) cooled to ∼ -20 °C with the Exos2 water cooling system (Koolance, Auburn, WA, USA). Images were acquired with HCImage software (Hamamatsu, Hamamatsu, Japan), binned to 4 × 4 and cropped to a 16 × 512-pixel region of interest, necessary to achieve the high image acquisition rates. Images were saved in CXD format.

Stimulation of APs was achieved with an extracellular tungsten parallel bipolar electrode (FHC, Bowdoin, ME, USA) mounted on a PatchStar motorized micro-manipulator (Scientifica, Uckfield, United Kingdom). To reliably induce an AP, we delivered a 1 ms 10 mV pulse at the soma. The timing of the LED illumination, the acquisition and stimulation were triggered externally through Clampex software (pClamp 10, Molecular Devices, San Jose, CA, USA).

All GEVI Imaging recordings were performed in presence of NBQX (10 µM), Gabazine (10 µM) and D-2-amino-5-phosphonovalerate (APV, 25 µM; all from Bio-Techne, Wiesbaden-Nordenstadt, Germany) in order to block synaptic transmission and ensure that the observed events were due to the stimulation only. IbTx was delivered using a custom gravity-fed perfusion system and bath volume was maintained by a peristaltic pump (Watson-Marlow 120s; Watson-Marlow, Rommerskirchen, Germany).

### Image analysis

Image analysis was performed as previously described by Sabater *et al.*^28^. In short, voltage imaging recordings were analyzed in ImageJ and custom scripts within MATLAB software as follows. Prior to analysis, if drift had occurred in the x and y axis during the recording, the images were aligned using the TurboReg ImageJ plugin^86^. Images were then imported into MATLAB and fluorescence intensity profiles were obtained from regions of interest (ROIs) manually drawn around neuronal structures identified visually on a maximum projection image of the time series by averaging across the ROI pixels in each frame. The fluorescence profile was separated into single trials and corrected for background camera noise by subtracting the average intensity value of the timepoints where no illumination light was applied. Then, recordings were reconstructed at a sampling rate of 100 kHz using cubic spline interpolation. Bleaching was estimated by fitting a single exponential function to each individual trace smoothed using an averaging filter with a window of 3 and interpolated. The resulting curve was used to correct an unfiltered interpolated version of the recording. An accurate representation of the AP waveform with an enhanced SNR was obtained by averaging over the repeats and extracting the AP parameters from the resulting trace. The fluorescence profile during the 20 ms preceding a response was used as baseline.

### Solutions and reagents

Reagents were purchased from Sigma-Aldrich (St. Louis, MO, USA) if not indicated otherwise.

#### Conditions I, II, VII and XI

The extracellular bath solution contained (in mM): 122 NaCl, 3 KCl, 10 D-Glucose, 26 NaHCO_3_, 1.25 Na_2_HPO_4_, 1.3 MgCl_2_, and 2.0 CaCl_2_. After equilibration with 95% O_2_–5% CO_2_, pH was adjusted to 7.35 if needed by NaOH or HCl. The intracellular pipette solution contained (in mM): 130 K gluconate, 1 MgCl_2_, 10 HEPES, 0.3 EGTA, 10 Tris-Phosphocreatine, 4 Mg-ATP, 0.3 Na-GTP. When testing the contribution of BK channels, 300 nM Iberiotoxin (Abcam, Cambridge, United Kingdom) was included in the bath solution.

#### Conditions III and IV

The extracellular bath solution contained (in mM): 125 NaCl, 3 KCl, 25 Glucose, 25 NaHCO_3_, 1.25 Na_2_HPO_4_, 1.1 MgCl_2_, and 1.1 CaCl_2_. After equilibration with 95% O_2_–5% CO_2_, pH was adjusted to 7.35 if needed by NaOH or HCl. The intracellular pipette solution contained (in mM): 150 K Gluconate, 10 K-HEPES, 3 Mg-ATP, 0.3 Na-GTP, 0.05 EGTA, 10 NaCl.

#### Conditions V and VI

The extracellular bath solution contained (in mM): 150 NaCl, 4 KCl, 10 HEPES, 10 glucose, 1.0 MgCl_2_, and 2.0 CaCl_2_, and was equilibrated with 95% O_2_–5% CO_2_. The intracellular pipette solution contained (in mM): 135 K gluconate, 4 NaCl, 0.05 EGTA, 10 HEPES, 3 MgCl_2_, 0.3 NaGTP, 3 Na_2_ATP, 10 Na_2_-Creatine phosphate with pH normalized to 7.25 using KOH.

#### Condition VIII

The extracellular bath solution contained (in mM): 122 NaCl, 3 KCl, 1.25 NaH_2_PO_4_, 26 NaHCO_3_, 10 glucose, 1.3 MgCl_2_, and 2.0 CaCl_2_, and was bubbled with 95% O_2_–5% CO_2_. The intracellular pipette solution contained (in mM): 130 K gluconate, 0.3 EGTA, 10 HEPES, 2 MgCl_2_, 0.3 NaGTP, 4 Na_2_ATP (or Mg-ATP), 10 Na_2_-Creatine phosphate (or Tris-Creatinine phosphate).

#### Conditions IX and X

The extracellular bath solution contained (in mM): 145 NaCl, 2.5 KCl, 10 HEPES, 24 glucose, 1.3 MgCl_2_, and 2.0 CaCl_2_, and was equilibrated with 95% O_2_–5% CO_2_. The intracellular pipette solution contained (in mM): 140 K gluconate, 2 NaCl, 0.5 EGTA, 20 HEPES, 3 MgCl_2_, 0.3 NaGTP, 4 Na_2_ATP, with pH adjusted to 7.25 using KOH.

#### Organotypic hippocampal neuron recordings

The external bath solution contained (in mM): 125 NaCl, 26 NaHCO_3_, 3 CaCl_2_, 2.5 KCl, 2 MgCl_2_, 0.8 NaH_2_PO_4_, and 10 D-glucose, and was equilibrated with 95% O_2_–5% CO_2_. The intracellular pipette solution contained (mM): 120 K gluconate, 20 KCl, 10 HEPES, 0.5 EGTA, 2 MgCl_2_, 2 Na_2_ATP, and 0.3 NaGTP (pH 7.4).

### Statistics

If not otherwise noted, data are expressed as box plots with boxes reflecting 25^th^–75^th^ percentile. The median is indicated by a line within the box, whiskers indicate 10^th^ and 90^th^ percentile. Sample sizes (n) provide the number of individual somatic recordings. All data sets were obtained from ≥ 2 independent cell cultures. Data in Fig. 3A were tested for statistical difference using Fisher’s exact test. Confidence limits for 95% confidence intervals were calculated as Wilson interval^87^. Other experimental groups were independent and tested by the Mann-Whitney *U* test. Tests were performed in IGOR PRO or GraphPad Prism 8. Results were considered significant at p < 0.05.

## Acknowledgment

This work was supported by a European Research Council Consolidator Grant (ERC CoG 865634) the German Research Foundation (HA6386/10-2) to S.H. and by grants awarded by the U.S. Department of Veterans Affairs (BX002547) and NIGMS (GM134110) to S.M.S.

## Declaration of interests

The authors declare no competing interests.

## Figure supplements

**Figure 2 – figure supplement 1.**
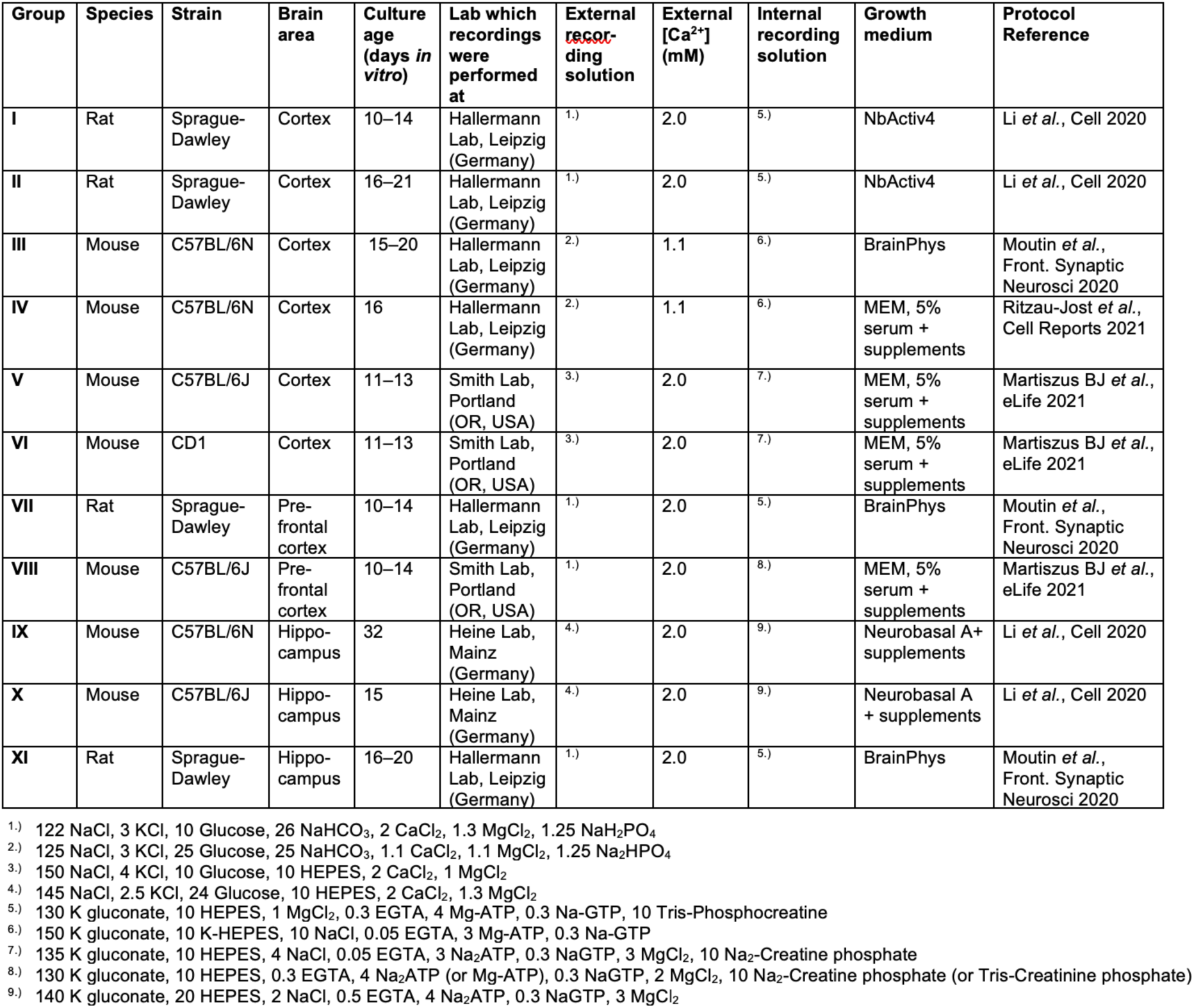
Additional information on recording conditions. The table lists additional relevant information on the experimental conditions I – XI.

**Figure 2 – figure supplement 2.**
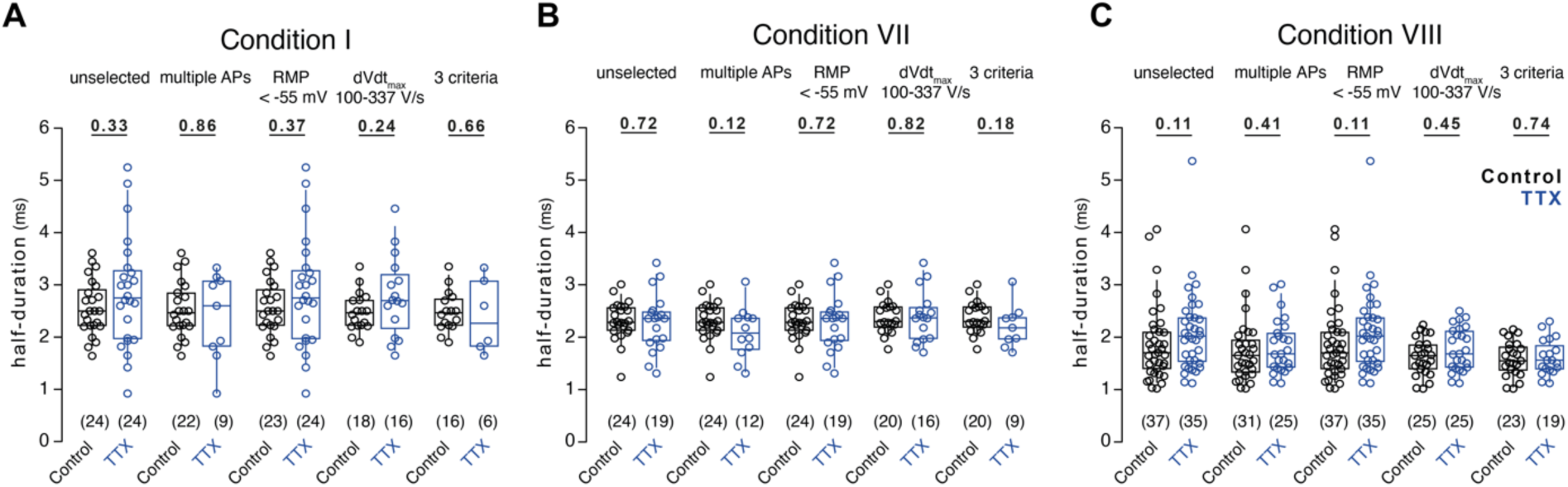
Absence of TTX-induced AP broadening in further selected conditions. (A) Box plots of AP durations under control condition (black) or upon TTX-treatment (blue) recorded in condition I either unselected or selected for parameters reflecting unimpaired neuronal function. (B) AP durations as in A for recordings in condition VII. (C) AP durations as in A for recordings in condition VIII. Numbers of recorded neurons provided in brackets. Box plots depict median ± interquartile range, whiskers reflect 10–90 percentile. P-values calculated by Mann-Whitney-*U* test.

**Figure 4 – figure supplement 1.**
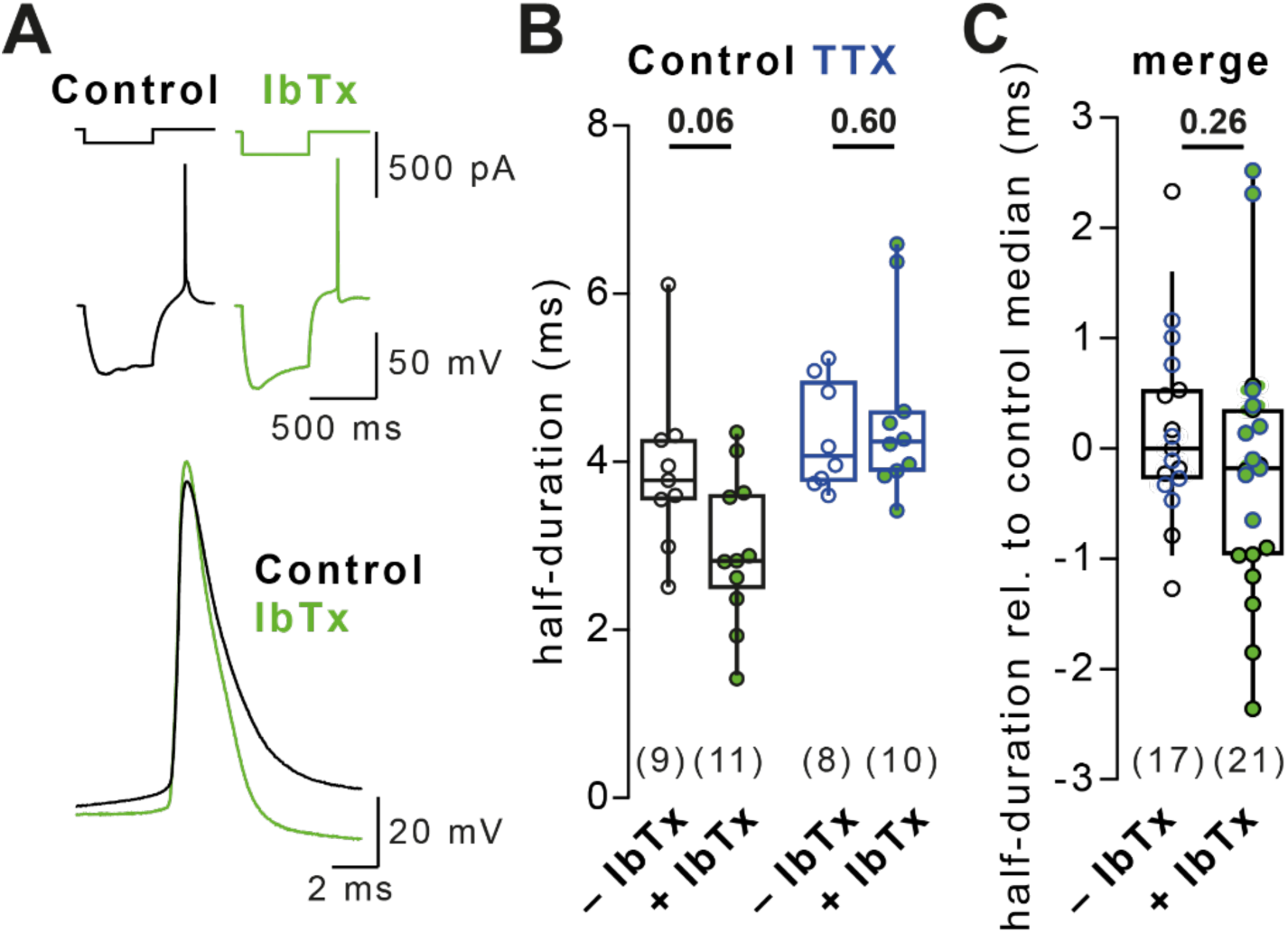
BK channel block does not affect rebound AP duration. (A) Example rebound APs in control solution (black) and solution containing 300 nM Iberiotoxin (IbTx, green). (B) AP duration in control and TTX-treated cells in IbTx-free (black and blue, respectively) or IbTx-containing solution (green). Data recorded in condition I. (C) Merged AP duration across control and TTX-treated cells for recordings in IbTx-free and IbTx-containing solution (data in A, normalized to the respective group’s median duration in IbTx-free solution). Numbers of recorded neurons provided in brackets. Box plots depict median ± interquartile range, whiskers reflect 10–90 percentile. P-values calculated by Mann-Whitney-*U* test.

## Notes

### Competing Interest Statement

The authors have declared no competing interest.

### Summary of Updates

We have made changes to the text and small changes to the figures to clarify raised questions.

